# Varying crosslinking motifs drive the mesoscale mechanics of actin-microtubule composites

**DOI:** 10.1101/554584

**Authors:** Shea N. Ricketts, Madison L. Francis, Leila Farhadi, Michael J. Rust, Moumita Das, Jennifer L. Ross, Rae M. Robertson-Anderson

## Abstract

The cytoskeleton dynamically tunes its mechanical properties by altering the interactions between semiflexible actin filaments, rigid microtubules, and crosslinking proteins. Here, we use optical tweezers microrheology and confocal microscopy to characterize how varying crosslinking motifs impact the microscopic and mesoscale mechanics and mobility of actin-microtubule composites. We show that, upon subtle changes in the crosslinking pattern, composites separate into two distinct classes of force response – primarily elastic versus more viscous behavior. For example, a composite in which actin and microtubules are crosslinked to each other is markedly more elastic than one in which both filaments are crosslinked but cannot link together. Notably, this distinction only emerges at mesoscopic scales in response to nonlinear forcing, whereas varying crosslinking motifs have little impact on the microscale mechanics and steady-state mobility of composites. Our unexpected scale-dependent results not only inform the physics underlying key cytoskeleton processes and structures, but, more generally, provide valuable perspective to materials engineering endeavors focused on polymer composites.

## Introduction

The multifunctional mechanics of eukaryotic cells are largely determined by the cytoskeleton, a dynamic network of interacting biopolymers and binding proteins. Two principal cytoskeletal biopolymers are actin filaments and microtubules. Actin filaments polymerize from globular actin monomers to ∼7 nm wide semiflexible filaments with a persistence length *l*_*p*_ ≈ 10 *μ*m^1,2^, while microtubules are formed by tubulin dimers polymerizing into 25 nm wide hollow rigid tubes with *l*_*p*_ ≈ 1 mm^2-6^. Networks of semiflexible actin filaments play critical roles in cell polarity and contractile and migratory processes^7,8^; while microtubules mediate intracellular trafficking and transport, chromosomal dynamics, and mitotic spindle alignment during cell division^7,9^. Synergistic interactions between these two filaments, which are mediated by steric and chemical interactions (i.e. entanglements and crosslinking)^1,8,10,11^, establish essential cell asymmetries and enable proliferation, differentiation, and migration^1,8,12-14^. While crosslinking of both filaments is ubiquitous in cells, serving important roles in locomotion, membrane reinforcement, and intercellular cargo transport^4,8,14^, there is mounting evidence of crosslinking *between* actin and microtubules, mediated by proteins such as tau, MAP2, APC, profilin, and plectin^8,15-21^. Specifically, actin-microtubule co-crosslinking has been shown to be important to cortical flow, wound healing, neuronal cone growth, cell migration, and muscle contraction^7-9,14,15,17,22-25^.

Aside from the obvious physiological relevance, investigations of actin-microtubule composites with variable crosslinking motifs are motivated by materials engineering, as they serve as a model platform for designing and investigating flexible-stiff polymer composites^2,11,26,27^. In order to increase stiffness, strength, and toughness in composite materials, soft polymer networks are often reinforced with stiffer, load bearing fibers such as nanotubes and carbon nanofibers^11,26,28,29^. Further, varying types of crosslinking have been used to reinforce and increase the mechanical strength of flexible-stiff polymer double-networks^30-32^. Crosslinking has also been shown to preserve memory and reduce network deformation in composite polymer hydrogels^32,33^.

Here, we use optical tweezers microrheology and confocal microscopy to characterize the mechanics and mobility of actin-microtubule composites with varying crosslinking motifs. We create equimolar composites of actin and microtubules in which: only actin is crosslinked (*Actin*), only microtubules are crosslinked (*Microtubule*), both actin and microtubules are crosslinked (*Both*), and actin and microtubules are crosslinked to each other (*Co-linked*) (Fig. 1a). We characterize mechanics by optically pulling a microsphere through the composites and simultaneously measuring the induced force during strain as well as the post-strain relaxation of force. We find that the mesoscale force response separates into two distinct classes, with one class exhibiting a nearly complete elastic response and enhanced post-strain mechano-memory, while the other class exhibits a softer more viscous response and substantial force dissipation following strain. We show that these two classes, which are only readily apparent at mesoscopic scales (>*μ*m), are dictated by the degree to which microtubules are crosslinked.

**Figure 1.**
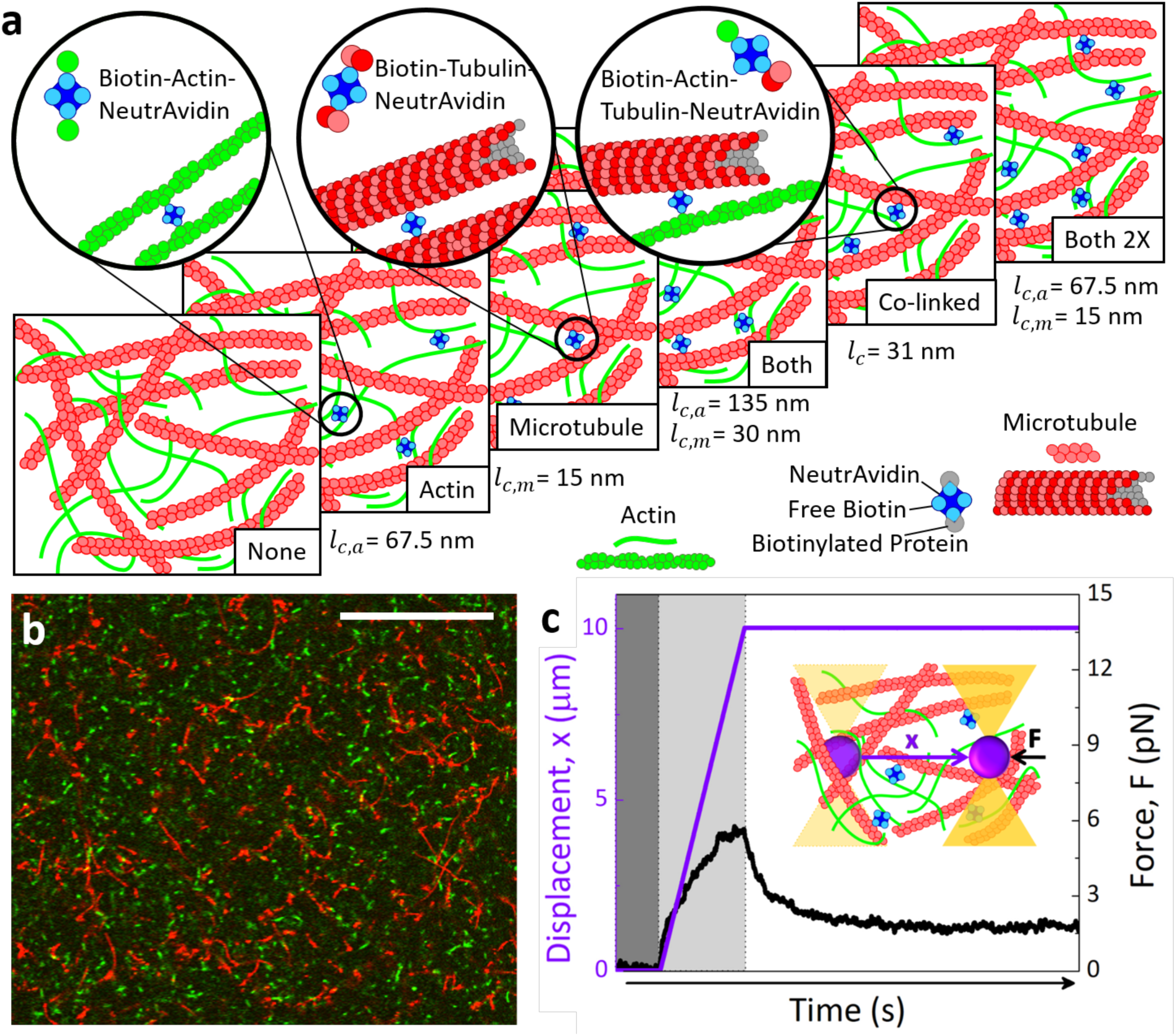
Schematic of experiments to probe the role that crosslinking plays in the mechanics and mobility of actin-microtubule composites. (**a)** Cartoon of molecular components comprising actin-microtubule composites with varying crosslinker motifs (*None, Actin, Microtubule, Both, Co-linked, Both 2x*) described in the text. The calculated length between crosslinkers for each motif is listed under each cartoon, where *l*_*c,a*_ is the length between crosslinkers along an actin filament, *l*_*c,m*_ is the length between crosslinkers along a microtubule, and *l*_*c*_ is the length between crosslinkers when actin and microtubules are linked to each other. (**b)** Two-color laser scanning confocal micrograph of 5.8 *μ*M *Co-linked* actin-microtubule composite with ∼3% of actin and microtubules labeled with Alexa-488 (green) and rhodamine (red), respectively. (**c)** Sample microrheology data for a *Co-linked* composite, showing the microsphere displacement (purple) and the force measured for an individual trial (black) before (dark grey region), during (light grey region) and after (white region) the microsphere displacement. Inset: Cartoon of optical tweezers microrheology measurement. An optically trapped 4.5 *μ*m diameter microsphere (not drawn to scale) embedded in a crosslinked composite is displaced 10 *μ*m (*x*, purple) at a speed of 10 *μ*m/s while the corresponding force the composite exerts is measured (*F*, black).

Despite the relevance of actin-microtubule composites to biology, physics, and engineering, most in vitro studies to date have focused on single-component networks of either actin or microtubules^4,34-48^. In response to strain, entangled actin networks have been reported to exhibit varying degrees of stiffening, softening, and yielding depending on the actin concentration and the scale of the strain^36,38,39^. Crosslinked actin networks exhibit enhanced stiffness and prolonged, more pronounced elasticity compared to entangled solutions^35,37,43,44^. Entangled actin networks have also been shown to relax force via an array of mechanisms including bending, stretching, retraction, reptation, and confinement-hopping^39,49-52^. While many of these mechanisms are suppressed by crosslinking, crosslinked networks have been reported to exhibit relaxation and plasticity due to crosslinker unbinding/rebinding, slippage, and stretching^53-55^. Compared to entangled actin networks, entangled microtubules have been reported to display enhanced elasticity and stiffening, as well as more suppressed relaxation^4,9,42^. Additionally, microtubule bending plays a more important role than stretching in the force relaxation^56^. While the timescales for microtubules to relax have been suggested to be on the order of several minutes or more^4,42^, crosslinked microtubule networks have been reported to display some degree of relaxation due to temporary force-induced crosslinker unbinding events^4^.

Few studies have investigated composite networks of actin and microtubules^11,27,57,58^ due, in part, to the incompatibility of standard polymerization conditions for each protein^27^. However, methods have recently been introduced to co-polymerize actin and tubulin in situ to create randomly-oriented co-entangled composites of actin and microtubules^27^. The first study that employed these methods used microrheology and microscopy techniques to show that a surprisingly large molar fraction of microtubules (>70%) was needed to substantially increase the response force of the composites. This intriguing result was attributed to the larger mesh size of microtubule networks compared to actin networks of the same molarity. Further, actin-rich composites softened in response to strain while microtubule-rich composites stiffened. This distinction arose from actin bending (leading to softening) that was increasingly suppressed by microtubules (leading to stiffening). This result corroborated a previous study that investigated the nonlinear macrorheology of crosslinked actin networks doped with low concentrations of microtubules, in which the high bending stiffness of microtubules served to suppress actin bending and enhance stretching in response to strain^11,43^.

## Results and Discussion

To systematically investigate the role that crosslinking plays in composites of actin and microtubules we design actin-microtubule composites with four distinct crosslinking motifs. We use our previously established protocols^27^ to create equimolar co-entangled actin-microtubule composites. We then incorporate biotin-NeutrAvidin complexes into composites to selectively crosslink the filamentous proteins, keeping the crosslinker:protein ratio fixed at *R* = 0.02^37^. We create composites with: crosslinked actin (*Actin*), crosslinked microtubules (*Microtubule*), crosslinkers equally forming actin-actin and microtubule-microtubule bonds (*Both*), and crosslinkers binding actin to microtubules (*Co-linked*) (Fig. 1a). We compare these composites to a composite without crosslinkers (*None*) as well as one in which both filaments are crosslinked but *R* is doubled (*Both 2x*) (Fig. 1a). While all composites have the same crosslinker density *R* and protein concentration, the length between crosslinkers along actin filaments *l*_*c,a*_, along microtubules *l*_*c,m*_, and along both filaments *l*_*c*_ (for *Co-linked*) vary for the different composites, as listed in Figure 1a and described in Methods.

### Mechanical Response Dynamics

To characterize the mechanical response of our designed composites we use optical tweezers microrheology to drag microspheres through each composite and measure the local force applied to each sphere by the composite during and following the strain. Figure 2a shows the measured force, *F(x)*, as a microsphere is pulled 10 *μ*m through each type of composite. As expected, all crosslinked composites exhibit overall larger resistive forces than the composite with no crosslinkers (*None*). The more interesting and unexpected result that emerges is that the force curves appear to display two distinct responses dependent on the crosslinking motif: pronounced softening behavior in which the slope of the force curves approach zero at large distances, versus largely elastic behavior in which *F(x)* remains linear with *x* until the end of the strain. To better delineate between these two behaviors, we normalize *F(x)* by the terminal force *F*_*t*_ reached when *x* = 10 *μ*m, which we notate as 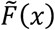. As shown in Figure 2b, 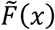 distinctly separates into two classes: Class 1 comprises *None, Actin*, and *Both* composites while Class 2 comprises *Microtubule, Co-linked*, and *Both 2x* composites.

**Figure 2.**
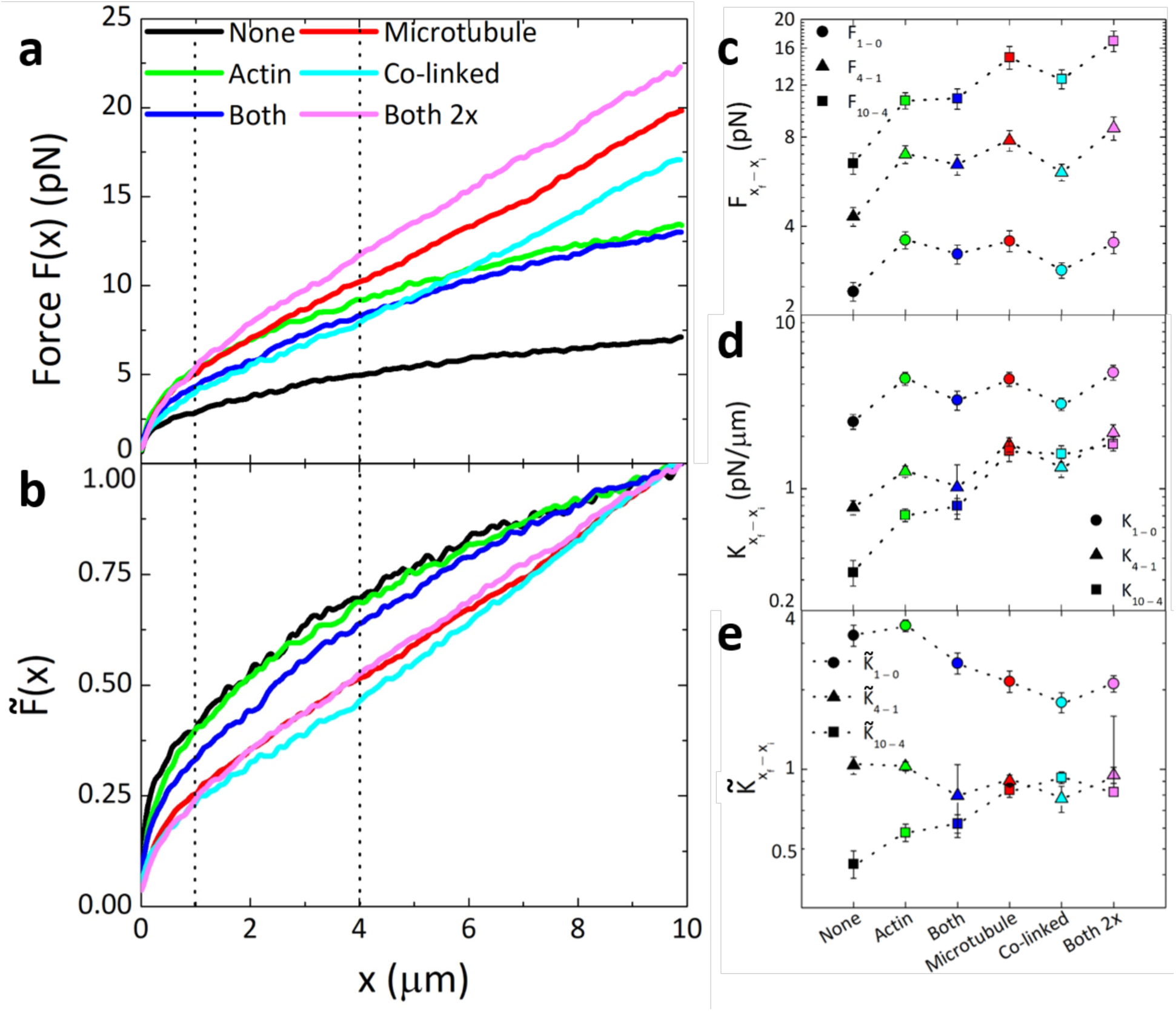
Mesoscale force response of crosslinked actin-microtubule composites display two distinct classes of behavior mediated by the degree of microtubule crosslinking. (**a**) Force *F(x)* that composites with varying crosslinking motifs (listed in legend) exert to resist the 10 μm strain. Dotted vertical lines separate the different regions over which *F(x)* and the stiffness *K* are averaged in (c)-(e). (**b**) The force shown in (a), normalized by the terminal force *F*_*t*_ reached at the end of the strain, denoted as 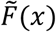. As shown, composites separate into two distinct classes – those that exhibit substantial softening and yielding (Class 1: *None, Actin, Both*) versus those that exhibit nearly complete elastic response (Class 2: *Microtubule, Co-linked, Both 2x*). (**c**) Force values from (a), averaged over different bead displacements *x*_*f*_ *– x*_*i*_, denoted as *F*_*xf–xi*_ (color coded as in (a)). Data shown is for *x*_*f*_ *– x*_*i*_ = 1 – 0 *μ*m (circles), 4 – 1 *μ*m (triangles) and 10 – 4 *μ*m (squares). (**d**) Stiffness, *K = dF/dx*, averaged over the same displacements as in (**c**), as listed in the legend. (**e)** The dimensionless stiffness, 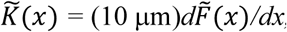, averaged over the same displacements as in (**c**). Error bars in (c)-(e) are calculated from computing the standard error of force and stiffness values measured for each individual trial (∼50-60 trials per data point).

For a purely elastic material we expect to see a linear relationship between the resistive force and distance the microsphere is pulled (*F ∼ x*), and once the microsphere stops the force the network exerts on the sphere *F(t)* should be maintained with no relaxation or dissipation (mechano-memory, *F(t) ∼ F*_*t*_). Conversely, a purely viscous material yields a strain-independent force response (*F ∼ x*^*0*^) that is reached as soon as the microsphere reaches constant speed (nearly immediately), and once the microsphere stops the network immediately and completely relaxes its force on the sphere (*F(t) ∼ 0*). Most biopolymer networks, including entangled actin-microtubule composites, display viscoelastic behavior in between these two extremes.^2,27^ However, crosslinking can lead to more elastic response by suppressing network rearrangements and polymer bending modes that can dissipate stress^11,37^. Figure 2b shows that Class 2 composites exhibit nearly complete elastic response while Class 1 exhibits softening and yielding to viscous behavior. The distinction between the two classes appears to be the degree to which microtubules are crosslinked, which we elaborate on further below.

Only composites in which microtubules are crosslinked (*Microtubule, Co-linked, Both 2x*) exhibit elastic response, while *Actin* and *None* force curves exhibit similar curvature indicative of more viscous response. This result suggests that at this crosslinking density, crosslinks do not substantially prevent semiflexible actin filaments from bending and/or rearranging to dissipate stress. Further, at a given *R* the length between crosslinks along an actin filament is ∼4*x* longer than that for microtubules (see Methods), so there are comparatively fewer crosslinks along an actin filament to suppress relaxation modes. The counterintuitive result presented in Figure 2 is that a composite in which both filaments are crosslinked (*Both*) exhibits soft/viscous behavior similar to *None* and *Actin* despite the fact that both actin and microtubules are crosslinked. Given the role that crosslinking plays in enhancing elasticity, one would expect that a network in which all filaments are crosslinked would be more elastic than one in which just half of the proteins are linked (*Actin* or *Microtubule*). However, because in this composite both actin and microtubules are crosslinked and the density of crosslinkers remains fixed (*R* = 0.02), there are half as many crosslinkers available to crosslink microtubules than for *Microtubule* and *Co-linked* composites. Thus, the length between crosslinkers along each filament is doubled.

To determine if this increased *l*_*c,m*_ and *l*_*c,a*_ is indeed the reason for more viscous behavior for *Both* composites, we perform experiments in which we double the crosslinker density (*Both 2x*) such that the length between crosslinkers along each actin filament and microtubule is the same as in *Actin* and *Microtubule* composites, rather than being twice as long. As shown in Figure 2, upon doubling *R*, the composite exhibits Class 2 elastic response similar to *Microtubule* and *Co-linked* networks. Thus, it appears that the critical requirement for elasticity is not only that microtubules must be crosslinked but that they must be crosslinked such that the length between crosslinkers is sufficiently small. Reducing *l*_*c,m*_ leads to a more connected network, thereby constraining filament fluctuations and rearrangements and leading to enhanced network stiffness^62^.

To quantify the distinct behaviors of Class 1 and Class 2 composites and examine the lengthscale dependence of the response, we average *F(x)* over different microsphere displacements *x*_*f*_ *– x*_*i*_, which we denote as *F*_*xf–xi*_ (Fig. 2c). As shown, at submicron distances (0 – 1 *μ*m) there is little difference between the force response the two classes exhibit, with all crosslinked composites displaying an average force of *F*_*1–0*_ ≈ 3 ± 0.6 pN. This result indicates that the varying crosslinking motifs have little effect on the microscale, linear regime response of composites. Even at somewhat larger microscale displacements (1 – 4 *μ*m), while crosslinked composites exhibit larger forces than *None*, the type of crosslinking has little effect on the measured force (with all values lying within ∼1 pN of each other). Only at mesoscale distances, on the order of the length of filaments and substantially larger than the mesh size of composites, does the dramatic effect of crosslinking motif emerge, with Class 1 composites exhibiting an average force of *F*_*10–4*_ ≈ 9 ± 3 pN while Class 2 composites display *F*_*10–4*_ ≈ 15 ± 2 pN.

This scale-dependent variation is likely a result of both variable length between crosslinkers and a small fraction of crosslinker unlinking/relinking events. At the submicron level, the differences in crosslinker lengths only result in small differences in the density of crosslinkers encountered by the microsphere, and filament bending and stretching is minimal. As the distance becomes orders of magnitude larger than the crosslinker lengths, the effect of varying crosslinking densities, as well as the varying contributions of bending and stretching, become more appreciable. Further, at larger displacements (i.e. longer times), crosslinks that were initially ruptured (contributing to submicron softening) will have had time to re-link. While new un-linking events can occur, the force response at lengthscales >>*l*_*c*_ reflects that of a steady-state unbinding/rebinding regime, and the response will still be dominated by stretching and/or bending modes. The varying degree to which Class 1 and Class 2 composites can undergo each mode manifests as a clear distinction between measured force values at these lengthscales.

The mechanisms described above would more clearly contribute to differences in the elasticity or stiffness of the networks rather than the absolute magnitudes of the force responses. In fact, as shown in Figure 2b, the clear distinction between Class 1 and Class 2 is in the lengthscale-dependent curvature, which quantifies the elasticity, rather than the magnitudes of the force response. To better quantify this distinct behavior, we calculate the stiffness *K* of composites, analogous to the differential modulus in bulk rheology, which we quantify as *K = dF/dx* (Fig. 2d). We also compute the dimensionless stiffness 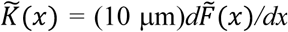, which quantifies the curvature displayed in Figure 2b (Fig. 2c). Larger *K* and 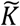 values are indicative of stiffer, more elastic networks while softer more viscous behavior results in smaller *K* and 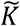 values.

As shown in Figure 2, both classes exhibit similar *K* values for submicron scales (*K*_*1–0*_ ≈ 4 ± 1 pN/*μ*m), and exhibit softening (decreasing *K*) from *K*_*1–0*_ to *K*_*4–1*_. Class 1 composites do soften slightly more than Class 2 composites over the first 4 *μ*m; however, the differing degrees of softening for Class 1 and Class 2 composites is much more substantial at larger lengthscales (>4 *μ*m). *K*_*10–4*_ values drop to ∼0.6 ± 0.2 pN/*μ*m for Class 1 composites, whereas for Class 2 composites *K*_*10–4*_ values are indistinguishable from *K*_*4–1*_ values (no softening). To better quantify the extent to which composites soften or remain elastic, we turn to 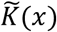. As shown, 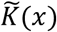 values for Class 1 composites drop by a factor of ∼3 from (0 – 1 *μ*m) to (1 – 4 *μ*m), and drop another ∼30% from (1 – 4 *μ*m) to (4 – 10 *μ*m). Class 2 composites, on the other hand, drop by a factor of ∼2 from (0 – 1 *μ*m) to (1 – 4 *μ*m) but remain constant for lengthscales beyond 4 *μ*m.

Our results and interpretations are validated by prior measurements on actin networks crosslinked by biotin-NeutrAvidin^37^. In this previous work, authors found that *R* > 0.05 was needed for actin networks to exhibit largely elastic response, with *R* < 0.05 networks displaying softening and yielding to a viscous regime over a 10 *μ*m bead displacement of comparable speed. The authors explained the transition to elastic response as arising from actin filaments stretching in the direction of the strain rather than bending non-affinely. This study also showed that force-induced crosslinker unbinding contributed to the yielding and relaxation behavior of low *R* networks. Such force-induced unbinding of biotin-NeutrAvidin crosslinkers has also previously been reported for crosslinked microtubule networks^4^.

Further, our results differ from previous actin studies in which the onset of elastic response was nearly immediate. Here, our Class 2 composites display softening behavior, similar to that of Class 1 until ∼2 *μ*m at which point a more elastic response takes over. This universal softening at small scales supports the presence of force-induced unbinding of both actin and microtubule crosslinks, as the length between crosslinkers is substantially smaller than a micron for all composites, whereas the deformation lengthscales needed to induce appreciable bending or stretching are much larger^35,37,59^. As the strain proceeds, the contributions from bending and stretching grow, outweighing that of crosslinker-unbinding and thus minimizing its effect on the force response. Further, unbinding events are reversible, and re-binding has been shown to contribute to sustained elastic response (coupled with partial dissipation) similar to that depicted by Class 2 composites (Fig. 2c-e)^4,35,37,59-61^.

### Relaxation Dynamics

As described above, elastic and viscous systems should display starkly different relaxation behaviors following strain, with purely elastic systems exhibiting no relaxation and viscous systems exhibiting immediate and complete force relaxation. Viscoelastic systems display behavior in between these two extremes, with viscoelastic biopolymer networks exhibiting both exponential and power-law relaxation over varying timescales^2,27,36-38,63^. To characterize the relaxation dynamics of the composites, we measure the time-dependent force *F(t)* exerted on the bead by each composite for 15 s following the strain (Figs. 1,3).

**Figure 3.**
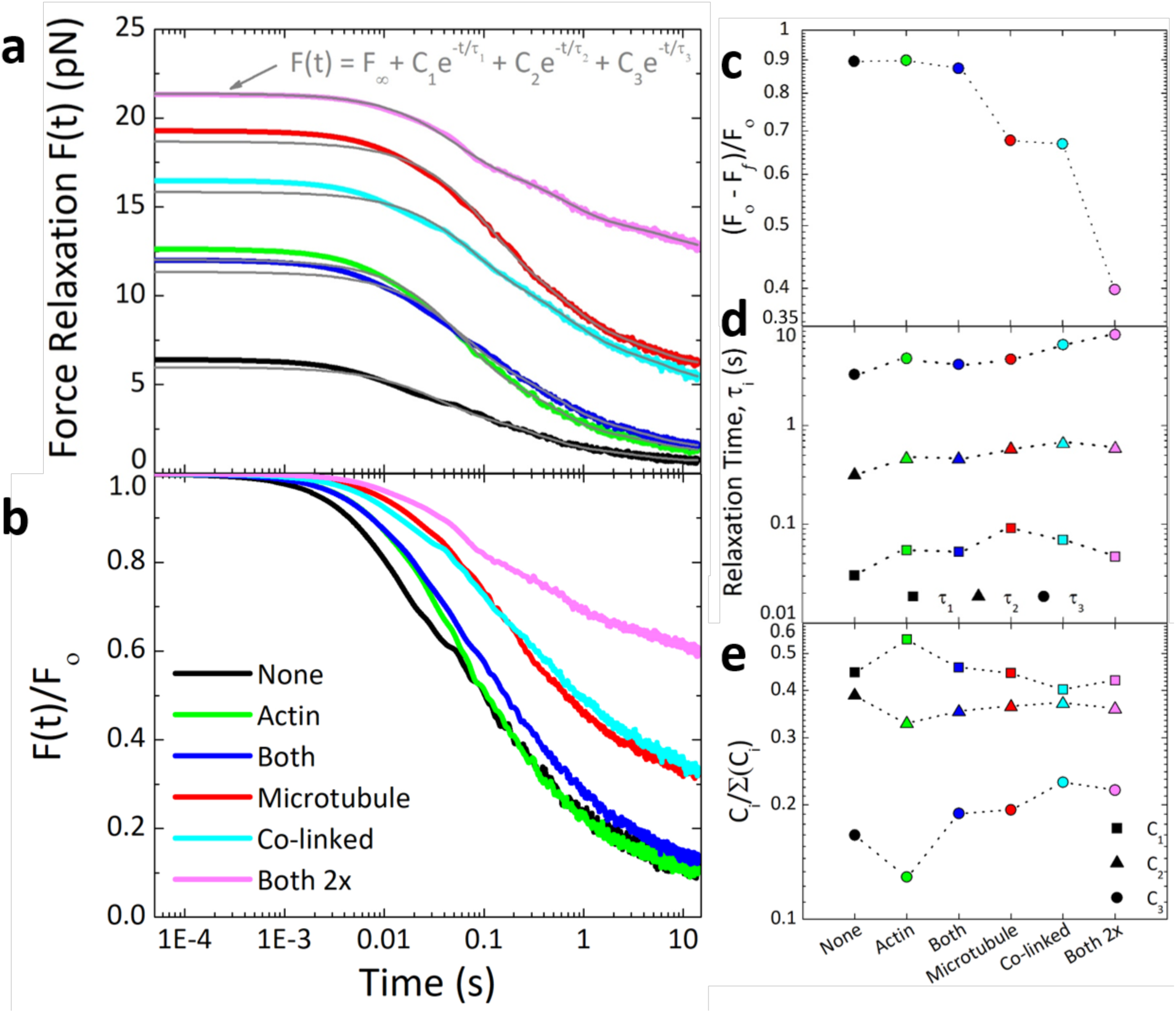
Actin-microtubule composites display multi-mode force relaxation with varying degrees of mechano-memory tuned by the crosslinker motif. (**a**) Force relaxation after microsphere displacement as a function of time *F(t)* for composites with varying crosslinking motifs listed in the legend in (b). Grey curves correspond to fits of each curve to the exponential sum equation displayed. (**b**) *F(t)* normalized by the force at the beginning of relaxation, *F*_*o*_. (**c**) The fractional force dissipation, computed as *(F*_*o*_ *– F*_*f*_*)/F*_*o*_ where *F*_*f*_ is the force at the end of the relaxation measurement period. As shown, Class 1 composites exhibit much more force relaxation compared to Class 2 composites that retain substantial memory of imposed strain. (**d**) Relaxation times, *τ*_*1*_ (squares), *τ*_*2*_ (triangles), and *τ*_*3*_ (squares) determined from the fits shown in (a). (**e**) Relative relaxation coefficients *C*_*i*_ */(C*_*1*_*+C*_*2*_*+C*_*3*_) for each exponential term (*i* = 1,2,3) determined from the fits to the data (shown in (a)).

As shown in Figure 3a, all composites exhibit some degree of force relaxation, though the extent to which the force decays depends on the crosslinking motif. This dependence becomes distinctly apparent upon normalizing *F(t)* by the initial force (immediately after strain) *F*_*0*_ (Fig. 3b). Class 1 composites exhibit faster and more complete relaxation than Class 2 composites which retain a substantial fraction of the initial force at the end of the measurement period. To quantify the varying degrees to which composites retain initial force, indicative of elastic behavior, we evaluate the fractional force dissipation *(F*_*o*_ *– F*_*f*_*)/F*_*o*_, in which *F*_*f*_ is the force reached at the end of the relaxation period. For reference, in a purely elastic network with no dissipation *(F*_*o*_ *– F*_*f*_*)/F*_*o*_ should be 0 whereas a viscous system should exhibit a value close to 1. As shown in Figure 3c, a clear difference between Class 1 and Class 2 composites emerges once again. Class 1 composites dissipate ∼89% of initial force whereas Class 2 composites only dissipate ∼40 – 68% of the initial force.

To determine the macromolecular mechanisms underlying the observed relaxation behavior, we evaluate fits of the force curves to a range of exponential and power-law functions that have been previously used to fit relaxation data for entangled and crosslinked biopolymer networks^27,36-38,48^. We find that the force relaxations for all composites can be well fit to a sum of three exponential decays with well-separated time constants (Fig. 3a). The function used for all composites, *F(t) = F*_*∞*_ *+ C*_*1*_*e*^*-t/τ1*^ *+ C*_*2*_*e*^*-t/τ2*^ *+ C*_*3*_*e*^*-t/τ3*^, includes an offset *F*_*∞*_ that is a measure of the amount of force that the composite retains indefinitely (*F*_*∞*_ ∼ *F*_*0*_ and 0 for elastic and viscous systems respectively). The three distinct decay constants, with similar values for all composites, indicate that three different relaxation mechanisms with well-separated characteristic timescales contribute to the force relaxation of all composite types (Fig. 3d). The decay times, averaged over all composite types, are *τ*_*1*_ = 0.06 ± 0.02 s, *τ*_*2*_ = 0.51 ± 0.12 s and *τ*_*3*_ = 5.28 ± 1.83 s. However, we note that all three timescales are ∼2*x* faster for the unlinked composite (*None*) compared to the crosslinked composites, which we discuss further in the next section. To evaluate the relative contribution of each mechanism to the force relaxation, we compare the corresponding fractional coefficients for each exponential term *C*_*i*_ */(C*_*1*_*+C*_*2*_*+C*_*3*_) (*i* = 1,2,3) (Fig. 3e). For all composites, the fast timescale contributes the most to the relaxation (∼40 – 54%), followed by the intermediate timescale (∼33 – 39%), and finally the slowest timescale (< 23%). However, Class 1 composites exhibit higher *C*_*1*_ values and lower *C*_*3*_ values than Class 2 composites, indicating that the slowest relaxation mechanism plays a more prominent role in the relaxation of Class 2 versus Class 1 composites.

To understand our fast relaxation timescale, we estimate the fastest predicted relaxation mode for entangled actin networks, which is termed the mesh time *τ*_*mesh*_ and can be thought of as the time for hydrodynamic effects to become important (i.e. for each filament to “feel” its neighbors)^39,64^. The mesh time for our composites, computed via *τ*_*m*_ *≈ βζξ*^4^*l*_*p*_^−1^ where is *β* = 1/*k*_*B*_*T, ζ* is the translational friction coefficient and *ξ* is the composite mesh size (see Methods)^39,63,64^, is ∼0.086 s, which is quite close to our fast relaxation time *τ*_*1*_ ≈ 0.06 s. Thus, we can interpret our fastest relaxation mechanism as arising from hydrodynamic effects of the surrounding mesh. This timescale relies on filaments being able to fluctuate and rearrange over lengthscales comparable to the mesh size. These fluctuations are hindered by rigid crosslinkers, so for smaller crosslinker lengths (Class 2) this relaxation mode should contribute less to the force relaxation. This effect can be seen in the varying *C*_*1*_ values for the different composites. Fast relaxation contributes to ∼48% of the relaxation for Class 1 compared to ∼42% for Class 2.

Another key mode of stress relaxation accessible to actin filaments and, to a lesser extent, microtubules is bending^3,65^. We estimate timescales for actin and microtubule bending using *τ*_*B*_ ≈ (*γ/κ*)[*L/*(3π/2)]^4^ where *κ ≈ l*_*p*_*k*_*B*_*T* is the bending rigidity, *L* is the filament length, *γ* = (4π*η*/ln(2*ξ/r*) is the perpendicular drag coefficient, *η* is the solvent viscosity, and *r* is the filament radius^3,39^. Using our measured lengths for actin and microtubules in composites (*L*_*A*_ ≈ 8.7 *μ*m and *L*_*M*_ ≈ 18.8 *μ*m)^27^ we compute bending relaxation times of *τ*_*B,a*_ ≈ 0.54 s and *τ*_*B,m*_ ≈ 0.15 s for actin and microtubules, respectively. These timescales are in close agreement with our measured *τ*_*2*_ values (∼0.3 – 0.6 s), suggesting that the second relaxation mode in our experiments is due to bending of both actin and microtubules. However, Class 1 composites display larger *τ*_*2*_ values than Class 2, with an average value of ∼0.6 s which is quite close to the predicted *τ*_*B,a*_. Conversely, the average *τ*_*2*_ value for Class 2 composites is ∼0.4, lying between *τ*_*B,a*_ and *τ*_*B,m*_. This distinction suggests that while microtubules can freely bend in response to large strains when they are not crosslinked or minimally crosslinked (Class 1), when *l*_*c,m*_ becomes sufficiently small (Class 2), microtubules can no longer bend to release stress. This suppression of bending modes leads to the elastic response seen in Figure 2 and the mechano-memory seen in Figure 3c.

To understand the slowest relaxation mode, which occurs on the order of ∼1 – 10 s for all composites, we turn to previous experiments investigating the stress relaxation of crosslinked actin networks following nonlinear strain^37^. These studies reported the presence of two distinct relaxation mechanisms available to actin filaments, which both occur on similar timescales as our measured *τ*_*3*_ and are both due to force-induced crosslinker unbinding and rebinding: curvilinear diffusion (reptation) and lateral confinement-hopping^37^. More specifically, crosslinked filaments can evade the confinement of the surrounding crosslinked mesh by crosslinks momentarily unlinking and thereby releasing the constraint they impose of the filaments, allowing for reptation and lateral confinement hopping. Such force-induced temporary unbinding events have been previously reported for both actin and microtubule networks crosslinked with biotin-NeutrAvidin^4,37^. Because fast timescale fluctuations (*τ*_*1*_) are more suppressed in Class 2 composites, we expect this slow mode to contribute more to the relaxation of Class 2 composites compared to Class 1. Figure 3e shows exactly this phenomenon: the average *C*_*3*_ value for Class 2 is ∼0.21 compared to ∼0.16 for Class 1.

### Steady-State Mobility

To determine the steady-state mobility of actin and microtubules in each composite type, we analyze time-series of dual-color confocal images, as described previously^27^ and in Methods. For each time-series we compute the average standard deviation of the intensity over time for all pixels <*δ*> and normalize by the corresponding average intensity of all pixels over time <*I*> (Fig. 4). Higher values of this mobility metric indicate a more mobile network of filaments that fluctuates more while lower values imply a more static network.

**Figure 4.**
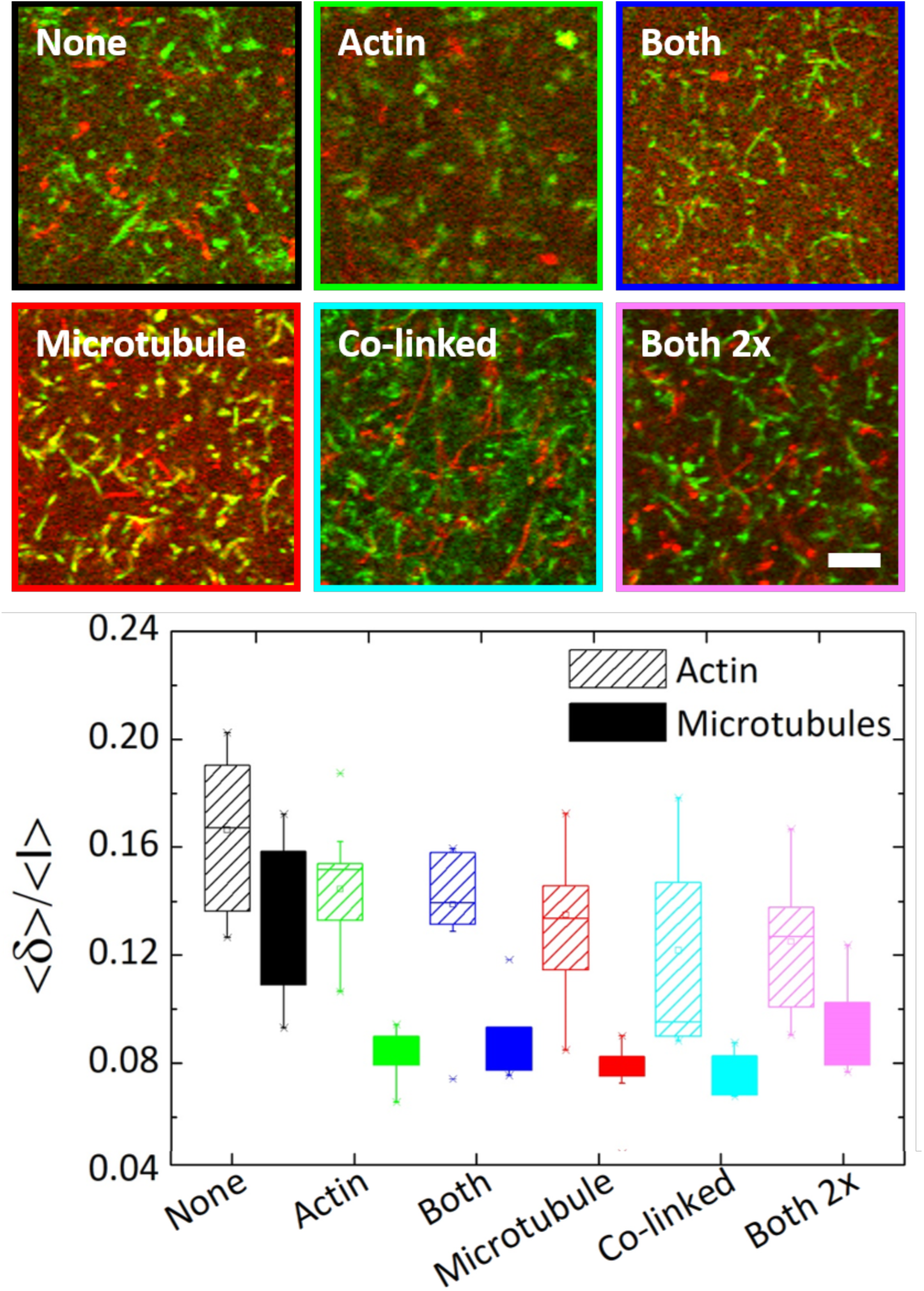
All crosslinking motifs reduce the steady-state mobility of actin and microtubules in composites. (Top) For each composite, a 128×128 image shows the standard deviation of intensity values for each pixel over time for actin (green) and microtubules (red) in a 60 s time-series. Scale bar is 10 *μ*m and applies to all images. (Bottom) Box-whisker plot of the steady-state mobility, determined by computing the average standard deviation of pixel intensities *<δ>* normalized by the overall average pixel intensity *<I>* for each time-series (as described in Methods). For each composite type, *<δ>/<I>* is calculated separately for actin and microtubules and data shown is for 10-12 time-series. As shown, microtubules are less mobile than actin filaments in all composites. Further, while crosslinking reduces the mobility of both filaments, the specific crosslinking motif has little effect.

Comparing *<δ>/<I>* values for actin and microtubules in the same composite, we find that actin filaments in all composites fluctuate significantly more than microtubules with an average *<δ>/<I>* value that is ∼1.5*x* greater than that for microtubules (*p* < 0.03 using KS-Test described in Methods, Fig. 4). This result, which is in line with our previous findings for composites without crosslinkers^27^, is not surprising given the ∼100-fold lower bending rigidity of actin compared to microtubules. Comparing the mobility of the crosslinked composites to the unlinked composite (*None*), we find that all crosslinking motifs reduce the mobility of both filaments by ∼1.4*x* (*p* ≤ 0.04 and *p* ≤ 0.02 for actin and microtubules, respectively). This result corroborates our relaxation data in which all three decay times for *None* are ∼1.85*x* faster than for crosslinked composites (Fig. 3d). Likewise, the measured resistive force during strain for *None* is substantially smaller than for crosslinked composites, indicating it is more fluid and mobile (Fig. 2).

However, while we see a clear distinction in the force response of Class 1 and Class 2 composites at mesoscopic scales (and in the nonlinear regime) (Fig. 2), the mobilities of both actin and microtubules display minimal variation between the different crosslinking classes (*p* > 0.09 for actin, *p* > 0.07 for microtubules). Because the mobility we are measuring is at the submicron scale (each pixel is 0.41 μm) and in the steady-state (linear) regime, we expect our mobility results to more closely match our microrheology results at the submicron and microscopic scales (corresponding to the low-force, linear regime). As described above, at these lengths and force scales there is little difference between the different crosslinking motifs, with all composites exhibiting similar force and stiffness values (Fig. 2). This effect is also seen in the relaxation data in which the relaxation times for all crosslinked composites are similar.

## Conclusion

Actin and microtubules form interacting networks within the cytoskeleton, providing cells with mechanical integrity and enabling a myriad of mechanical processes such as locomotion, morphogenesis, intracellular transportation, and division. Many of these diverse functions are mediated by crosslinking proteins that can bind actin, microtubules, or both proteins. Further, the role of crosslinking *between* actin and microtubules has received much recent attention and is thought to be responsible for maintaining cell shape and polarity, growth and structure of neurons, and spindle positioning in mitosis^8,15,16^. The design and characterization of composites of flexible and stiff polymers is also a topic of current interest in materials engineering^26,66^, yet how crosslinking can tune the mechanical response of these composites has remained largely unexplored.

Here, we characterize the mobility and mechanics – from submicron to mesoscopic scales – of custom-designed composites of actin and microtubules with distinct crosslinking patterns. We create composites in which actin is crosslinked (*Actin*), microtubules are crosslinked (*Microtubule*), both actin and microtubules are crosslinked (*Both*), and actin and microtubules are crosslinked to each other (*Co-linked*). We use optical tweezers microrheology and dual-color fluorescence confocal microscopy to measure the nonlinear force response and steady-state mobility of each composite type. We show that composites separate into two distinct classes which are dictated by the degree to which microtubules in each composite are crosslinked. Class 1, which comprises *None, Actin*, and *Both* composites, displays largely viscoelastic response to the imposed strain with substantial softening. Contrastingly, Class 2, comprised of *Microtubule, Co-linked* and *Both 2x* composites, elicit a nearly elastic response to perturbation. Notably, this distinction only emerges at mesoscopic scales in response to nonlinear forcing. Conversely, the crosslinking motif has little impact on the microscopic linear regime mechanics and steady-state filament mobility. These intriguing scale-dependent dynamics likely play a significant role in the ability of the composite cytoskeleton to mediate a vast array of different mechanical processes, and could be exploited to design multifunctional soft materials with scale-dependent mechanical properties. The custom-designed composites and measurement techniques demonstrated in this study further lay the groundwork for building complexity into in vitro cytoskeleton networks to determine how each component plays a role in cellular mechanics and in the design of novel biomimetic materials.

## Methods

Rabbit skeletal actin, biotinylated actin and Alexa-488-labeled actin were purchased from Cytoskeleton (AKL99, AB07) and Thermofisher (A12373) and stored at −80 °C in a Ca buffer (2 mM Tris (pH 8.0), 0.2 mM ATP, 0.5 mM DTT, 0.1 mM CaCl_2_). Porcine brain tubulin, biotinylated tubulin, and rhodamine-labeled tubulin were purchased from Cytoskeleton (T240, T33P, TL590M) and stored at −80 °C in PEM-100 (100 mM PIPES (pH 6.8), 2 mM MgCl_2_, and 2 mM EGTA).

Crosslinked networks were formed using preassembled biotin-NeutrAvidin complexes optimized previously for stable, isotropic crosslinking of actin networks^37^. As depicted in Figure 1a, crosslinker complexes are comprised of NeutrAvidin flanked on either side by biotinylated actin and/or tubulin, depending on the desired crosslinking motif. We incorporated crosslinkers into previously established protocols to form co-entangled actin-microtubule composites^27^. In short, actin monomers, tubulin dimers, and crosslinker complexes were added to PEM-100 supplemented with 1 mM ATP, 1 mM GTP, and 5 *μ*M Taxol. To image composites, 0.13 *μ*M of Alexa-488-labeled actin filaments and rhodamine-labeled microtubules, as well as oxygen scavenging agents [4.5 mg/ml glucose, 0.5% β-mercaptoethanol, 4.3 mg/ml glucose oxidase, 0.7 mg/ml catalase] were added (Fig. 1b). To enable microrheology measurements, carboxylated microspheres (Polysciences, Inc.), with a 4.5 *μ*m diameter, were also added (Fig. 1c). The solution was then mixed, pipetted into a ∼20 *μ*L sample chamber, and allowed to incubate at 37°C for 30 minutes to co-polymerize both proteins.

We prepared six different composite types that varied in their crosslinking motif as follows (Fig 1a): (1) no crosslinkers are present (*None*); (2) actin filaments in the composite are crosslinked to each other (*Actin*); (3) microtubules in the composite are crosslinked to each other (*Microtubule*); (4) both actin and microtubules are crosslinked but they cannot crosslink to each other (*Both*); (5) actin and microtubules are crosslinked to each other but not to themselves (*Co-linked*); and (6) a composite similar to *Both* but in which the densities of actin-actin and microtubule-microtubule crosslinks are both doubled (*Both 2x*).

For all presented data, total protein concentration was held fixed at 5.8 *μ*M with an equimolar ratio of actin to tubulin, and the molar ratio of crosslinker to protein was fixed at *R* = 0.02. The mesh size of actin and microtubule networks comprising each composite are *ξ*_*A*_ = 0.3/*c*_*A*_ ^1/2^ = 0.85 *μ*m and *ξ*_*M*_ = 0.89/*c*_*T*_ ^1/2^ = 1.58 *μ*m in which *c*_*A*_ and *c*_*T*_ are the actin and tubulin concentrations in units of mg/ml^34,42,67^. From these mesh sizes we calculated the effective composite mesh as *ξ*_*C*_ = (*ξ*_*M*_^*-3*^ *+ ξ*_*A*_^*-3*^)^−1/3^ = 0.81 *μ*m^27^. We estimated the length between crosslinkers for all three types of crosslinking used in composites (actin crosslinking, microtubule crosslinking, and actin-microtubule co-linking) as follows (Fig. 1a). The length between crosslinkers along an actin filament was determined by *l*_*c,a*_ = *½l*_*mon*_ × *R*^*−1*^ where *l*_*mon*_ = 2.7 nm is the length that each actin monomer adds to an actin filament. The ½ prefactor takes into account that each crosslinker is shared between two filaments. Similarly, the length between crosslinkers along a microtubule was calculated by *l*_*c,m*_ = *½*(*l*_*ring*_*/13)* × *R*^*−1*^ where every 13 tubulin dimers adds *l*_*ring*_ = 7.8 nm in length to the microtubule. For *Actin* and *Microtubule* composites, *l*_*c,a*_ = 67.5 nm and *l*_*c,m*_ = 15 nm. In the composite where crosslinkers are equally distributed between actin-actin links and microtubule-microtubule links (*Both*), *R* remains fixed so there are half as many crosslinkers available to each network. Thus, *l*_*c,a*_ and *l*_*c,m*_ both double (i.e. *l*_*c,a*_ = 135 nm, *l*_*c,m*_ = 30 nm). For composites in which both networks are linked but *R* is doubled (*Both 2x*) *l*_*c,a*_ and *l*_*c,m*_ remain the same as in the *Actin* and *Microtubule* cases. For *Co-linked* composites, the length between crosslinkers that link actin to microtubules was computed as *l*_*c,a-m*_ *= (l*_*c,a*_*l*_*c,m*_)^*1/2*^ = 31 nm.

The optical trap used in all microrheology measurements, which has been previously described and validated^27,36,37,68^, was formed by a IX71 fluorescence microscope (Olympus) outfitted with a 1064 nm Nd:YAG fiber laser (Manlight) and focused with a 60x 1.4 NA objective (Olympus). A position-sensing detector (Pacific Silicon Sensor) measured the deflection of the trapping laser, which is proportional to the force acting on the trapped microsphere. Stokes drag method was used to calibrate trap stiffness^69,70^. A nanopositioning piezoelectric stage (Mad City Labs) was used to move the trapped microsphere relative the sample (Fig. 1c).

For the data presented in Figures 1-3, a trapped bead was displaced 10 *μ*m at a constant speed of 10 *μ*m/s relative to the sample chamber. Stage position and laser deflection were recorded at 20 kHz during and after microsphere displacement using custom-written Labview codes. Custom-written MATLAB scripts were used for post-measurement analysis. Force curves displayed in Figures 1-3 are averages over 50 – 60 different trials taken between two samples. All trials were taken in different regions of the sample chamber, each separated by >100 *μ*m.

A Nikon A1R laser scanning confocal microscope with a 60x objective and QImaging QICAM CCD camera were used to collect 2D time-series of composites. The microscope is outfitted with 488 nm and 561 nm lasers and simultaneously records separate images for each laser channel (green and red) to separately visualize Alexa-488-actin (green) and rhodamine-tubulin (red) (Figs. 1b, 4). Time-series of 512×512 images (0.41 *μ*m/pixel) in each channel were recorded at 16 fps for 60 seconds. For each composite type, two different samples were imaged with 3-5 time-series taken for each sample.

Time-series from each channel (green and red) were analyzed separately to determine the mobility of actin and microtubules in the composites, as previously described^27^. In brief, every 16 frames in the time-series were averaged together to create time-series with 1 second frames. From these averaged time-series, we used FIJI/ImageJ to create a single collapsed image for each channel that represented the standard deviation of each pixel over time (Fig. 4). The average standard deviation over all pixels was then calculated to give a single standard deviation value for the entire time-series *<δ>*. This value, which represents the variation in intensity values for each pixel over time, is a measure of the extent to which filaments in the network fluctuate over time. To account for differences in overall intensity among different time-series, we also computed the average intensity over all pixels and frames for each time-series *<I>*, and normalized each standard deviation by this value. The resulting metric *<δ>/<I>*, which quantifies the mobility of actin and microtubules in each time-series, is presented in Figure 4. We carried out this analysis for all time-series collected for each composite type. We used the Kolmogorov-Smirnov statistical test (KS-Test, http://www.physics.csbsju.edu/stats/KS-test.html) to compare different data sets. From KS-tests, we computed probability values *p* for finding overlapping data and use the accepted value of *p* < 0.05 as the cutoff for determining when two data sets were significantly different.

## Author Contributions

S.N.R. designed and conducted microscopy and optical tweezer experiments, analyzed and interpreted data, and wrote the manuscript; M.L.F. conducted optical tweezer experiments and analyzed data; L.F. analyzed and interpreted microscopy data; M.J.R. helped analyze and interpret data; M.D. helped interpret data and write the manuscript; J.L.R analyzed and interpreted data and helped write the manuscript; R.M.R.-A. designed and advised experiments, analyzed and interpreted data, and wrote the manuscript.

### Acknowledgements

This research was funded by a National Science Foundation CAREER Award (grant no. 1255446, awarded to R.M.R.-A.); a National Institutes of Health R15 Award (National Institute of General Medical Sciences Award No. R15GM123420, awarded to R.M.R.-A.); and a William M. Keck Foundation Research Grant (awarded to R.M.R.-A., J.L.R., M.D., and M.J.R.)

